# Development of a rapid LC-MS/MS method for the simultaneous quantification of various flavonoids and phytohormones extracted from *Medicago Truncatula* leaves

**DOI:** 10.1101/2021.04.14.439919

**Authors:** Neema Bisht, Arunima Gupta, Pallavi Awasthi, Atul Goel, Divya Chandran, Neha Sharma, Nirpendra Singh

## Abstract

Flavonoids are small metabolites of plants, which are involved in the regulation of plant development as well as defence against pathogens. Quantitation of these flavonoids in plant samples is highly important and essential in the food and herbal industry. Hence, robust, reliable and sensitive methods are required for the analysis of these compounds in plant samples. In the present study, a high performance liquid chromatographic-tandem mass spectrometric (HPLC– ESI-MS/MS) method was developed and validated for the determination of nine flavonoids, including Liquiritigenin, Naringenin, Genistein, Daidzein, Formononetin, Biochanin A, 2’-hydroxy formononetin, 2’-methoxyformononetin, Medicarpin, and two phytohormones, Salicylic acid and Jasmonic acid, in *Medicago truncatula*. The analytes were separated by means of C-18, reversed-phase chromatography and detected using QTRAP mass spectrometer. Molecules were quantified using different transitions in positive and negative ion modes simultaneously in 12 minutes. The on-column limit of detection of all the analytes was as low as 0.03 pg, whereas the limit of quantification of all the compounds was observed upto 0.1 pg levels. Further the method was also validated in terms of specificity, linearity (r^2^> 0.99), average recovery (90.6 – 110.3%), accuracy (RE% ≤ 3%) and precision (RSD% ≤ 3%). As a proof of concept, the developed method was successfully used for the quantitation of these flavonoids and phytohormones from leaf extract of *Medicago truncatula*.

## 1. Introduction

Flavonoids, a widely-distributed class of polyphenol compounds in plants, have been studied for their various biological properties such as anti-cancer [1, 2], antioxidant [3, 4], anti-thrombotic vasodilatory [5], anti-allergic [6], and anticholinesterase [7], etc. These polyphenols are still extensively being explored for their role in plant metabolism, plant protection, and microbial interactions. Apart from their myriad functions in plants, flavonoids have received a great deal of attention due to their health promoting attributes such as anti-inflammatory, radical scavenging, antimicrobial, enzymatic inhibition, neuroprotective and phyto-estrogenic properties [8–11]. They are widely being used in the form of herbal formulations, herbal drinks, and supplements for their high antioxidant capacity [12].

Flavonoids comprise of more than 10,000 structures where the type and amount vary between plant species, different parts of the plant, different developmental stages and growth conditions. The structural diversity of flavonoids pose a great challenge in their analysis. Many analytical techniques have been employed [13, 14] to detect, structurally characterize [15] and quantify [16, 17] different classes of flavonoids for their critical evaluation in a wide range of matrices, including commercial products, edible parts of plants and biological fluids [14–17]. Till date, numerous UV, HPLC and LC-MS based methods for the determination and quantification of various flavonoids and phenolic acids have been described [18–29] but simultaneous identification and quantification of Daidzein, Liquiritigenin, Formononetin, Naringenin, Biochanin A, 2’-hydroxy formononetin, 2’-methoxy formononetin, Salicylic acid, Jasmonic acid, Medicarpin and Genistein has not been reported so far. The earlier studies have identified and quantified not more than seven flavonoids from the set for which the present method has been developed and validated [30, 31]. Also, the previously reported methods have certain limitations in term of reproducibility, linearity, limit of detection, limit of quantitation, inter-day and intra-day variability. The present study therefore attempts to quantitatively measure these metabolites in a sensitive and reproducible method in the leaves of *Medicago truncatula*.

## 2. Materials and methods

### 2.1 Reagents and chemicals

All chemicals and reagents used in this study were of ultrapure mass spectrometry grade. LCMS grade acetonitrile, water and methanol were procured from J.T baker (U.S.A) for mobile phase and sample preparation. Analytical grade formic acid from Sigma Aldrich (U.S.A) was used in the mobile phase. Reference standards for the flavonoids (Naringenin, Liquiritigenin, Genistein, Daidzein, Formononetin and Biochanin A) and phytohormones (Salicylic acid and Jasmonic acid) were procured from Sigma-Aldrich, USA.The flavonoids 2’-hydroxyformononetin, 2’-methoxyformononetin and Medicarpin were synthesized and purified in >98 % purities as described in the literature [32].

### 2.2 Instrumentation

#### 2.2.1 UHPLC

UHPLC experiments were performed on binary Exion UHPLC consisting of an autosampler, a thermostatic column oven and online degasser (Sciex USA). For chromatographic separations, Waters Acquity BEH C18 (1.7um, 2.1×50 mm; Security Guard ULTRA Cartridges, UPLC C18, 2.1mm) column was employed. Separations were achieved with mobile phase consisting of a mixture of 0.1% formic acid in water [A] and acetonitrile [B] in gradient elution mode at a flow rate of 0.2 ml/min. Mobile phase B was linearly increased from 20% to 80% in 10 min. Gradient was put on hold for 0.5 min and then again decreased to 20% of mobile phase B. Gradient was kept on hold again at 20% of B for 1 min. Total run time for the LC-MS/MS method was 12 minutes. The column oven was maintained at 40°C.

#### 2.2.2 Mass Spectrometry

Experiments were performed on QTRAP 6500+ mass spectrometer (Sciex, U.S.A), equipped with an ESI source and multi-component ion drive. Analyst™ 1.6.3 was used for data acquisition and parameter optimization. For quantitation, multiple reaction monitoring (MRM) analysis was used and quantitative analysis was performed on MultiQuant ™ software 3.0.2. The quadrupoles were operated at unit resolution. Out of the total eleven metabolites, seven metabolites were detected in the positive mode and four analytes were detected in the negative mode. The curtain gas, nebulizer gas and auxiliary gases were optimized at 35 L/min, 55 L/min and 60 L/min, respectively. Flavonoids were ionized using the Ion spray voltage of 5500 kV in the positive mode and 4500 kV in the negative mode, source temperature was kept at 550 °C, CAD gas was set to medium and mass acquisition range was set between 50 – 350 Da to get good quality MS spectra. The compound parameters were optimized using syringe infusion experiments **(Table 1)**. Transition reactions of the analytes are given in **Table 2**.

**Table 1:** Compound parameters.

| S.No | Analyte | Ionization mode | Parent ion (Q1) | Product ion (Q3) | DP | CE | CXP | EP |
| --- | --- | --- | --- | --- | --- | --- | --- | --- |
| 1. | Daidzein | Positive | 255 | 199 | 116 | 33 | 6 | 10 |
| 2. | Liquiritigenin | Positive | 257 | 137 | 56 | 29 | 10 | 10 |
| 3. | Formononetin | Positive | 269 | 197 | 91 | 49 | 12 | 10 |
| 4. | Naringenin | Positive | 273 | 153 | 31 | 31 | 10 | 10 |
| 5. | Biochanin A | Positive | 285 | 213 | 106 | 49 | 14 | 10 |
| 6. | 2'-hydroxy formononetin | Positive | 285 | 225 | 106 | 37 | 14 | 10 |
| 7. | 2'-methoxy formononetin | Positive | 299 | 148 | 111 | 39 | 10 | 10 |
| 8. | Salicylic acid | Negative | 136.9 | 65 | -35 | -38 | -17 | -10 |
| 9. | Jasmonic acid | Negative | 208.9 | 59 | -30 | -26 | -15 | -10 |
| 10. | Medicarpin | Negative | 269 | 254 | -31 | -43 | -10 | -10 |
| 11. | Genistein | Negative | 269 | 133 | -100 | -30 | -15 | -10 |

**DP:**Declustering potential; **CE:** Collision energy; **CXP:** Collision cell exit potential; **EP:** Entrance potential

**Table 2:** LOD & LOQ.

| Analyte | Retention Time (in min) | LOD (pg) | LOQ (pg) | Regression equation (y = mx + b) | Correlation coefficient |
| --- | --- | --- | --- | --- | --- |
| Daidzein | 2.9 | 0.10 | 0.36 | $y = 1053.12410x + (-)1048.69159$ | 0.997 |
| Liquiritigenin | 3.1 | 0.10 | 0.18 | $y = 3926.57344x + 1484.75979$ | 0.998 |
| Formononetin | 4.7 | 0.03 | 0.10 | $y = 3243.34734x + 13245.81315$ | 0.998 |
| Naringenin | 3.9 | 0.10 | 0.36 | $y = 9377.68417x + 9029.80333$ | 0.999 |
| Biochanin A | 5.9 | 0.18 | 0.36 | $y = 1051.33922x + (-)562.11882$ | 0.991 |
| 2'-hydroxy formononetin | 4.1 | 0.18 | 0.36 | $y = 6217.07537x + 18535.39120$ | 0.998 |
| 2'-methoxy formononetin | 4.5 | 0.06 | 0.10 | $y = 2798.60783x + 11840.05003$ | 0.998 |
| Salicylic acid | 3.0 | 0.36 | 0.72 | $y = 8810.71079x + 9.93007e4$ | 0.993 |
| Jasmonic acid | 4.1 | 0.10 | 0.72 | $y = 3651.53907x + 3226.30017$ | 0.999 |
| Medicarpin | 4.1 | 0.06 | 0.10 | $y = 830.29506x + 6696.81721$ | 0.998 |
| Genistein | 3.9 | 0.36 | 1.44 | $y = 1281.96120x + (-)10400.00791$ | 0.995 |

**Calibration curve range: 31.27ng/ml -1000ng/ml (31.27ppb - 1000ppb)**

### 2.3 Preparation of reference, standard and quality control solutions

All the purified flavonoids/reference standards were dissolved in methanol and stock solutions of 20ng/ml of each reference standard were prepared. Stock solutions were further diluted in methanol to obtain the working concentration of 1000 ng/ml. Working solution of 1000 ng/ml was serially diluted (dilution factor; 1:2) in methanol to obtain different concentrations of 500 ng/ml, 250 ng/ml, 125 ng/ml, 62.5 ng/ml, 31.75 ng/ml, respectively. 3μL of each of these reference standards was used to generate six calibration points.

Three quality control (QC) standards: LQC (Lower quality control) concentration, MQC (Middle quality control) concentration and HQC (Higher quality control) concentration were prepared in the concentration of 40 ng/ml, 80 ng/ml and 160 ng/ml respectively. Each of the standards were prepared for intra-day as well as inter-day analysis to check for precision and accuracy of the developed method. Another set of three different QC standards; LQC (50 ng/ml), MQC (100 ng/ml) and HQC (200 ng/ml) were used for recovery studies.

### 2.4 Extraction of flavonoids and phytohormones from plant samples

Metabolite extraction from the plant sample was performed as per Liu et al. (2017) [21] with slight modifications. Approximately 100 mg of *M. truncatula* leaf tissue was ground in liquid nitrogen and extracted three times in 1 ml of 80% HPLC-grade methanol (Merck, Germany). Each time the mixture was sonicated for 15 min at room temperature and centrifuged for 10 min at 15,000 ×g. In plants, flavonoids and phytohormones exist as free and conjugated forms like β-glucosides or acetyl and malonyl β -glucosides, which tend to hydrolyse in response to stimulus. To quantify the total concentration of these metabolites, plant extracts were treated with β - glucosidase to hydrolyse the conjugated forms of the flavonoids and phytohormones into their aglycone forms [33–36]. The supernatant was dried using a vacuum centrifuge and then incubated with 31.25 units/ml β-glucosidase (Sigma-Aldrich) at 37°C for 10-12 hours followed by vacuum drying. The pellet was dissolved in 100 μl of 20% methanol and a 1:10 dilution was filtered through a 0.20-μm polytetrafluoroethylene filter (Millipore, U.S.A).

## 3. Method validation procedures

### 3.1 Specificity and selectivity

Specificity and selectivity was established by comparing chromatogram of blank mobile phase with the chromatogram of spiked standards in different dilutions **(Fig. S2 & S3)**. The parent and daughter ion spectra were matched with the retention time of the reference materials to confirm the identity of the product ions and analytes.

### 3.2 Accuracy and precision

The precision and accuracy of the method were established in LQC, MQC, and HQC samples (6 replicates each in 3 sets) on the same day and on 3 consecutive days. The inter-assay and intra-assay accuracy (% RE) of the method was ascertained by calculating the absolute error from the difference of true value & measured value, with respect to true value as:

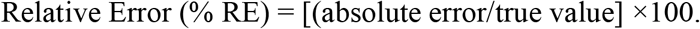

The intra-assay and inter-assay precision (% relative standard deviation or RSD) of the method was calculated from mean measured concentrations as:

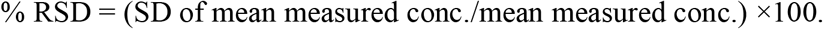

### 3.3 Linearity

To evaluate the linear relationship of the method, five different concentrations of standard flavonoids and phytohormones were prepared in methanol. The calibration curves were established and fitted by least square regression. The LOQ was defined as lowest concentration on the calibration curve with acceptable precision and accuracy (six replicates with RSD below 3% RE with in ±3%).

### 3.4 Stability and recovery

All the flavonoids and phytohormones are sensitive to temperature and oxidation. To prevent possible degradation under influence of temperature and environmental oxidation, all the stock solutions were stored in deep freezer (−20°C) and diluted at the time of analysis. Recovery studies were also performed by spiking eleven standards in the plant extract at three different concentrations, six replicates each for ensuring the optimum performance of the method.

## 4 Results and discussion

Flavonoids, the metabolites of plants play crucial functions in lifecycle of plants. Moreover, they also possess numerous health benefits for humans. The myriad functions played by flavonoids for plants as well as humans necessitates the development of rapid and reliable methods for their detection. Out of the several available methods, currently practiced mass spectrometry coupled with high performance liquid chromatography is considered to be the most versatile analytical technique in terms of data quality, chromatographic resolution and increased detection limits with greater sensitivity. In the present study, LCMS/MS was employed to optimize the simultaneous separation, identification and quantification of nine flavonoids and two phytohormones **(Fig. 1)**.

**Fig. 1:**
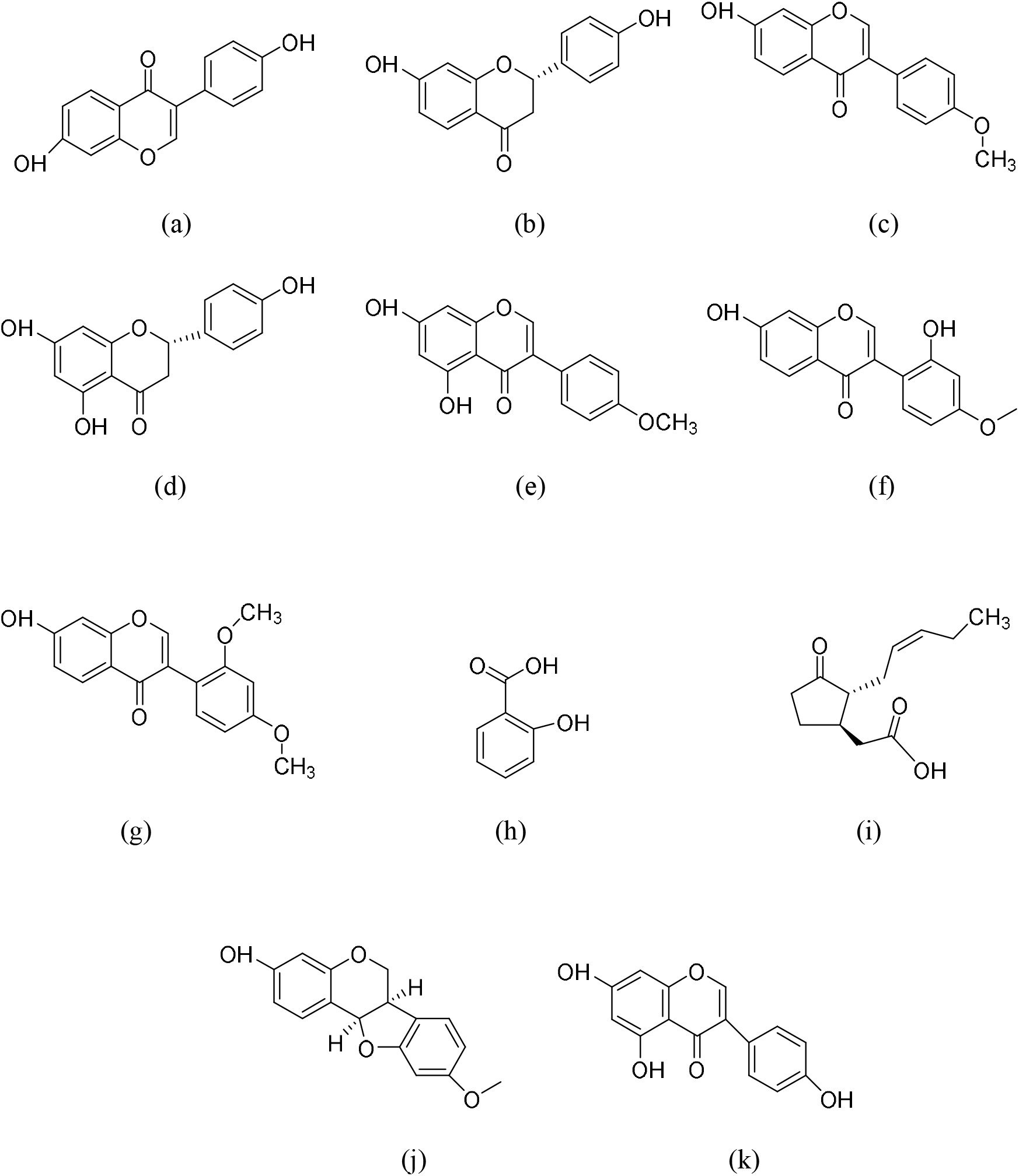
Chemical structures of flavonoids and phytohormones quantified in *Medicago truncatula* (a) Daidzein, (b) Liquiritigenin, (c) Formononetin, (d) Naringenin, (e) Biochanin A (f) 2’-hydroxyformononetin/Xenognosin, (g) 2’-methoxyformononetin, (h) Salicylic acid, (i) Jasmonic acid, (j) Medicarpin and (k) Genistein

### 4.1 Optimization of LC-ESI-Q-TRAP-MS parameters

Optimum chromatographic separation of eleven flavonoids and phytohormones from *Medicago truncatula* was achieved on Waters Acquity BEH C18 (1.7 um, 2.1×50 mm) column using our developed method. Gradient elution method allowed the separation and identification of all these compounds. The structures of these analytes are represented in **Fig. 1**. The resulting chromatograms show the respective peaks of the compounds at different retention times **(Table 2)**. Compound parameters in mass spectrometer consist mostly of voltages in the ion path. Optimal values for compound-dependent parameters vary depending on the compound being analyzed. Compound parameters such as declustering potential (DP), collision energy (CE), collision cell exit potential (CXP), entrance potential (EP) along with product ion (Q3) for individual standards were optimized using the Compound Optimization wizard in Analyst™ 1.6.3 version of Sciex mass spectrometer data acquisition software. In brief, a master mix of 100 ppb of all the eleven standards was prepared in 10% water/methanol and introduced into MS via direct infusion mode/ flow injection analysis (FIA) at the flow rate of 0.2 ml/min. The results of the product ion for individual standards were verified and confirmed by running MS2 scan for individual molecule. Quantification was done on the basis of single most abundant transition for each metabolite, including the ones co-eluting at similar retention times. The most abundant product ion was considered for quantification of each analyte.

### 4.2 LC-ESI-Q-TRAP-MS analysis

Optimum chromatographic separation of daidzein, liquiritigenin, formononetin, naringenin, biochanin A, 2’-hydroxy formononetin, 2’-methoxyformononetin, salicylic acid, jasmonic acid, medicarpin and genistein was achieved by acetonitrile: 0.1% formic acid in water with gradient elution. All the chemical analytes were identified based on the MS/MS ion simultaneously and the resulting chromatograms show a retention time of 2.9, 3.1, 4.7, 3.9, 5.9, 4.1, 4.5, 3.0, 4.1, 4.1, 3.9 min for daidzein, liquiritigenin, formononetin, naringenin, biochanin A, 2’-hydroxy formononetin, 2’-methoxyformononetin, salicylic acid, jasmonic acid, medicarpin and genistein, respectively **(Fig. 2A: MRM in positive ion mode & MRM in negative ion mode)**. A full scan in positive ion mode and negative ion mode was used for all the metabolites. During direct infusion, the mass spectra of the major product ions in the positive mode for daidzein, liquiritigenin, formononetin, naringenin, biochanin A, 2’-hydroxy formononetin, 2’-methoxyformononetin and in the negative mode for salicylic acid, jasmonic acid, medicarpin and genistein were used to monitor the ions of the compounds for selective MRM based quantification. The ions measured for the compounds were daidzein at m/z 255[M+H]^+^, liquiritigenin at m/z 257[M+H]^+^, formononetin at m/z 269[M+H]^+^, naringenin at m/z 273[M+H]^+^, biochanin A at m/z 285[M+H]^+^, 2’-hydroxy formononetin at m/z 285[M+H]^+^, 2’-methoxyformononetin at m/z 299[M+H]^+^, salicylic acid at m/z 136.9[M−H]^−^, jasmonic acid at m/z 208.9[M−H]^−^, medicarpin at m/z 269[M−H]^−^ and genistein atm/z 269[M−H]^−^. The single most abundant transitions used for quantification have been enlisted in **Table 1**. The fragmentation patterns for the transitions used have been represented in **Table 6**.

**Fig. 2A:**
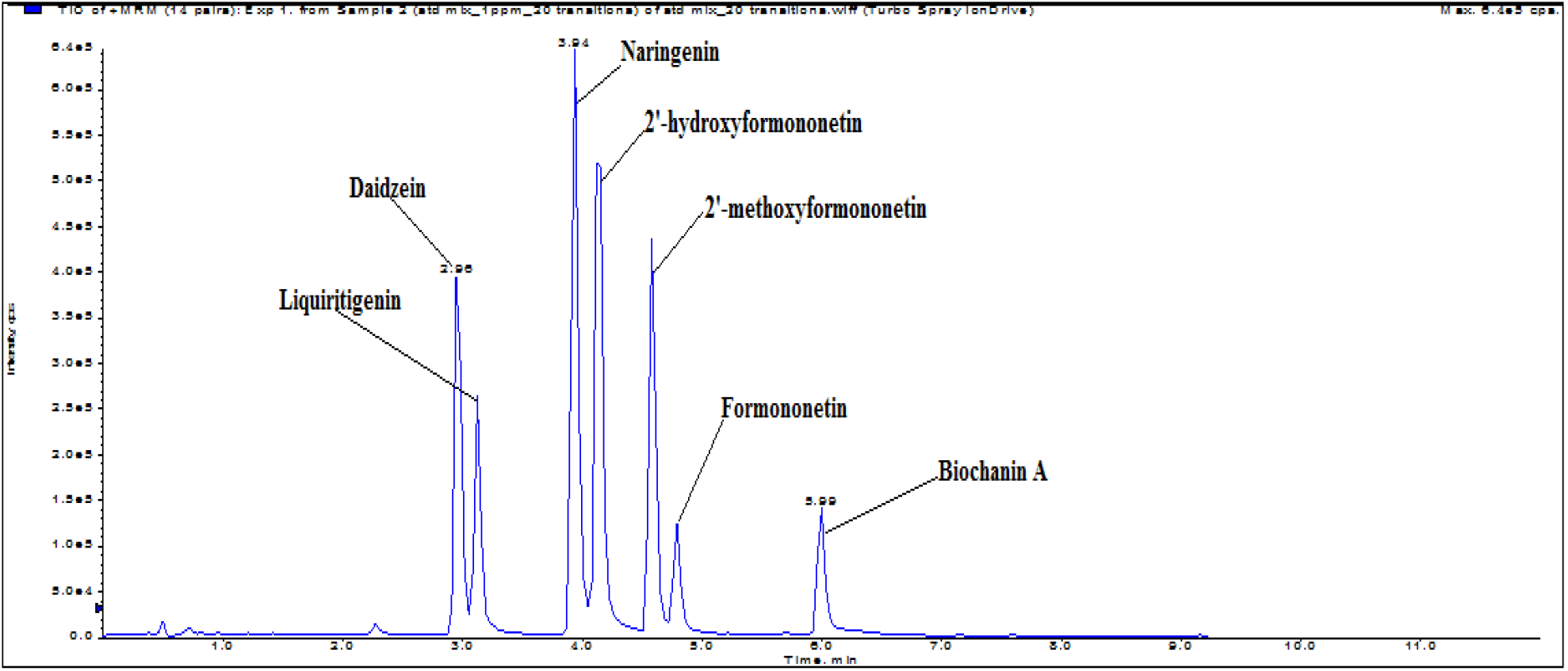
TIC of MRM scan in the positive mode

**Table 6:**
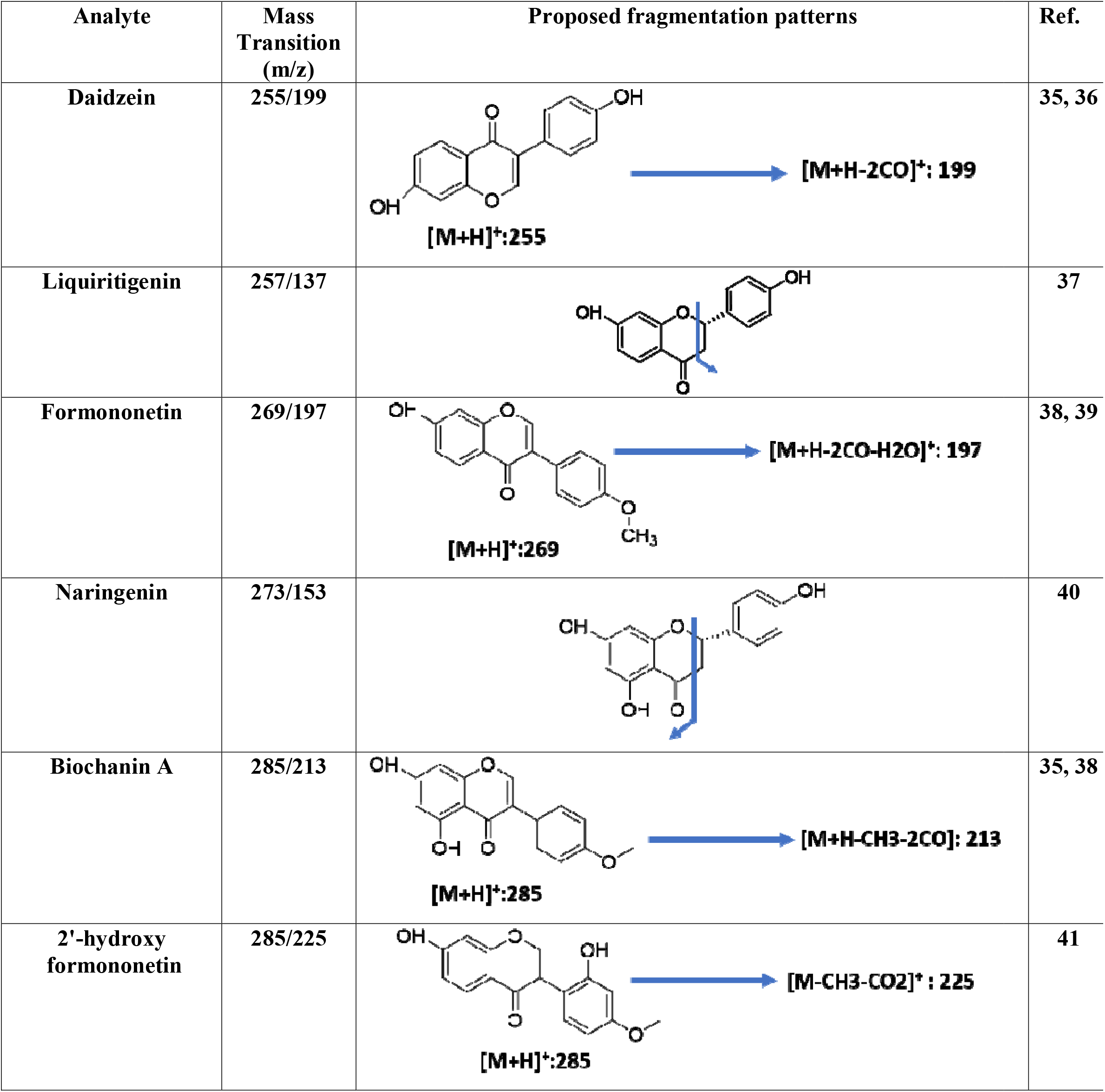
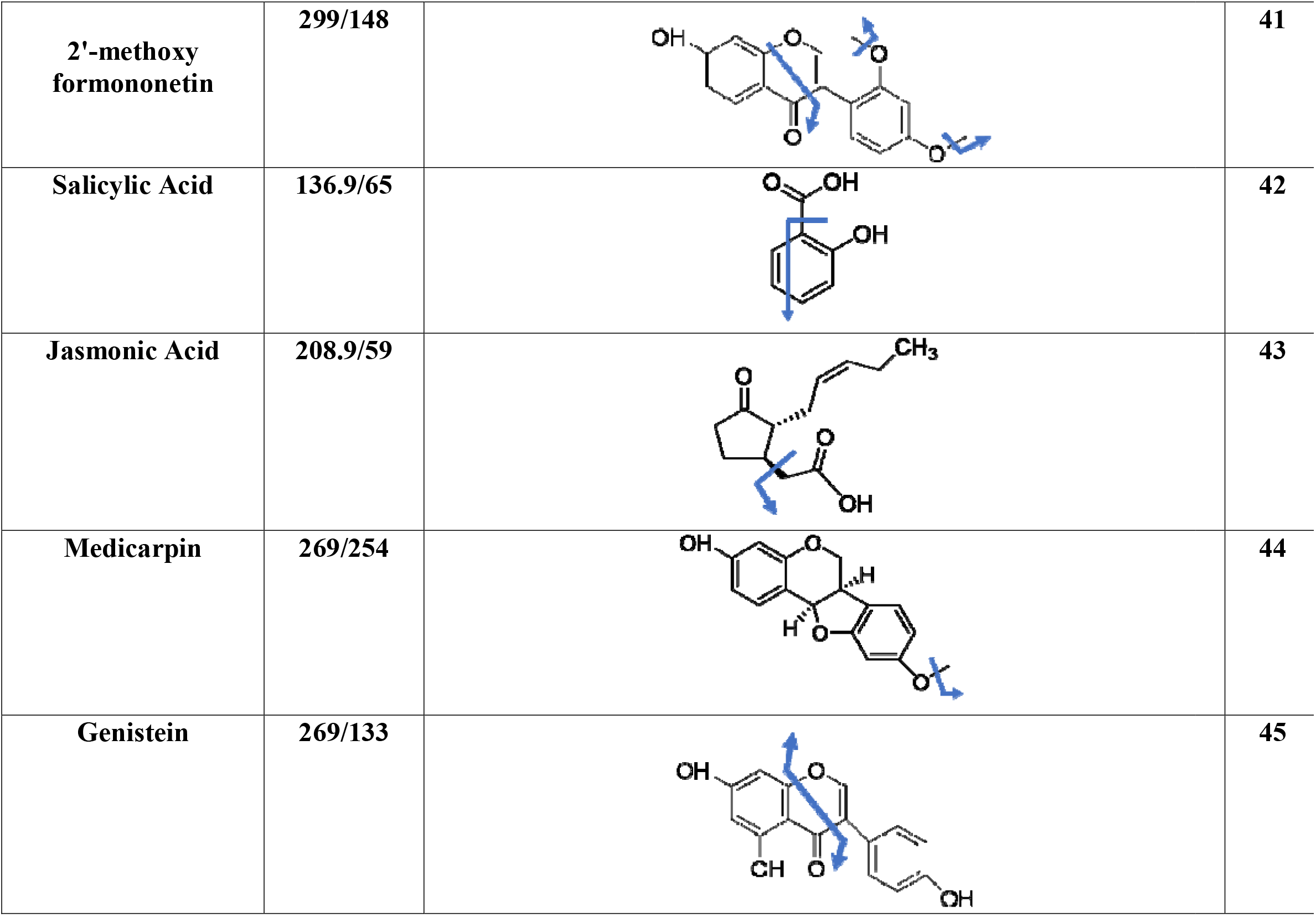
Fragmentation patterns of parent ion and transitions used for quantitation.

| Analyte | Mass Transition (m/z) | Proposed fragmentation patterns | Ref. |
| --- | --- | --- | --- |
| Daidzein | 255/199 | **[M+H]<sup>+</sup>:255** → **[M+H-2CO]<sup>+</sup>: 199** | 35, 36 |
| Liquiritigenin | 257/137 |  | 37 |
| Formononetin | 269/197 | **[M+H]<sup>+</sup>:269** → **[M+H-2CO-H2O]<sup>+</sup>: 197** | 38, 39 |
| Naringenin | 273/153 |  | 40 |
| Biochanin A | 285/213 | **[M+H]<sup>+</sup>:285** → **[M+H-CH3-2CO]: 213** | 35, 38 |
| 2'-hydroxy formononetin | 285/225 | **[M+H]<sup>+</sup>:285** → **[M-CH3-CO2]<sup>+</sup>: 225** | 41 |
| 2'-methoxy<br>formononetin | 299/148 |  | 41 |
| Salicylic Acid | 136.9/65 |  | 42 |
| Jasmonic Acid | 208.9/59 |  | 43 |
| Medicarpin | 269/254 |  | 44 |
| Genistein | 269/133 |  | 45 |

### 4.3 Method validation

The current method was validated in terms of its specificity, linearity, range, limit of detection, limit of quantification, recovery, accuracy and precision [37].

#### 4.3.1 Specificity and selectivity

The method possessed excellent non-interfering ability due to selective target ion validation approach (MRM). The parent and daughter ion spectra were matched with the retention time of the chromatogram of the reference materials to rule out any mismatch. The method was found to be specific, as mobile phase when compared with mobile phase spiked with standards did not show any interference at the respective retention times of any metabolite.

No carryover was observed from preceding sample into the next sample **(Supplementary data; Fig. S2 and S3).** The present method was specific and selective for the eleven metabolites in study.

#### 4.3.2 Linearity and range

The calibration curves for daidzein, liquiritigenin, formononetin, naringenin, biochanin A, 2’-hydroxy formononetin, 2’-methoxyformononetin, salicylic acid, jasmonic acid, medicarpin and genistein were linear over the concentration range of 31.75ng/ml to 1000ng/ml (r> 0.99). Results are shown in **Table 2. (Supplementary data; Fig. S1)**

#### 4.3.3 Limit of detection (LOD) and Limit of quantitation (LOQ)

The high sensitivity of the method enabled the detection of molecules upto picogram levels with formononetin being detected at 0.03 pg. The LOD and LOQ for daidzein, liquiritigenin, formononetin, naringenin, biochanin A, 2’-hydroxy formononetin, 2’-methoxyformononetin, salicylic acid, jasmonic acid, medicarpin and genistein are tabulated in **Table 2**.

#### 4.3.4 Recovery, Accuracy and Precision

The combined recovery for daidzein, liquiritigenin, formononetin, naringenin, biochanin A, 2’-hydroxy formononetin, 2’-methoxyformononetin, salicylic acid, jasmonic acid, medicarpin and genistein was carried out in LQC, MQC and HQC samples. The % recovery of daidzein was 90.6(from LQC), 92.2(from MQC), and 97.6(from HQC). The recovery % of liquiritigenin was 105.0(from LQC), 101.1(from MQC), and 101.1(from HQC). For formononetin, it was 100.7(from LQC), 102.0(from MQC), 101.4(from HQC). For naringenin, it was 105.0(from LQC), 106.2(from MQC), 102.5(from HQC). For biochanin A:106.7(from LQC), 106.5(from MQC), 103.0(from HQC). For 2’-hydroxy formononetin, it was 110.3(from LQC), 110.3(from MQC), 106.6(from HQC). For 2’-methoxyformononetin: 108.2(from LQC), 108.6(from MQC), 104.0(from HQC). For Salicylic acid, it was found to be 108.0(from LQC), 104.1(from MQC), 103.1(from HQC). For Jasmonic acid:106.5(from LQC), 100.9(from MQC), 101.7(from HQC). For medicarpin, it was 101.6(from LQC), 105.6(from MQC), 15.5(from HQC) and for genistein, the recovery % was found to be 96.8(from LQC), 97.6(from MQC), 98.7(from HQC). **(Table 3).**

**Table 3:** Recovery studies.

|  | Concentration<br>in Unspiked<br>sample<br>(ng/ml) | Spiked<br>concentration<br>(ng/ml) | Expected<br>concentration<br>(ng/ml) | Detected<br>concentration<br>(ng/ml) | Recovery% |
| --- | --- | --- | --- | --- | --- |
| Daidzein | 51.5 | 50 | 101.5 | 92 | 90.6 |
|  |  | 100 | 151.5 | 139.8 | 92.2 |
|  |  | 200 | 251.5 | 245.5 | 97.6 |
| Liquiritigenin | 487.3 | 50 | 537.3 | 564.2 | 105.0 |
|  |  | 100 | 587.3 | 594.1 | 101.1 |
|  |  | 200 | 687.3 | 695.2 | 101.1 |
| Formononetin | 5343.3 | 50 | 5393.3 | 5432 | 100.7 |
|  |  | 100 | 5443.3 | 5553 | 102.0 |
|  |  | 200 | 5543.3 | 5621 | 101.4 |
| Naringenin | 176.6 | 50 | 226.6 | 238 | 105.0 |
|  |  | 100 | 276.6 | 294 | 106.2 |
|  |  | 200 | 376.6 | 386.3 | 102.5 |
| Biochanin A | 29.6 | 50 | 79.6 | 85 | 106.7 |
|  |  | 100 | 129.6 | 138.1 | 106.5 |
|  |  | 200 | 229.6 | 236.5 | 103.0 |
| 2'-hydroxy<br>formononetin | 35.6 | 50 | 85.6 | 94.5 | 110.3 |
|  |  | 100 | 135.6 | 149.7 | 110.3 |
|  |  | 200 | 235.6 | 251.2 | 106.6 |
| 2'-methoxy<br>formononetin | 53.9 | 50 | 103.9 | 112.5 | 108.2 |
|  |  | 100 | 153.9 | 167.2 | 108.6 |
|  |  | 200 | 253.9 | 264.2 | 104.0 |
| Salicylic acid | 109.6 | 50 | 159.6 | 172.4 | 108.0 |
|  |  | 100 | 209.6 | 218.4 | 104.1 |
|  |  | 200 | 309.6 | 319.2 | 103.1 |
| Jasmonic acid | 156 | 50 | 206 | 219.4 | 106.5 |
|  |  | 100 | 256 | 258.5 | 100.9 |
|  |  | 200 | 356 | 362.4 | 101.7 |
| Medicarpin | 74.7 | 50 | 124.7 | 126.7 | 101.6 |
|  |  | 100 | 174.7 | 184.6 | 105.6 |
|  |  | 200 | 274.7 | 289.9 | 105.5 |
| Genistein | 132.6 | 50 | 182.6 | 176.8 | 96.8 |
|  |  | 100 | 232.6 | 227.2 | 97.6 |
|  |  | 200 | 332.6 | 328.5 | 98.7 |

The intra-assay accuracy in terms of % RE was in the range of 0.25 to 1.83 for daidzein,0.12 to 2.0 for liquiritigenin, −2.16 to −0.33 for formononetin, 1.08 to 1.33 for naringenin, 0.33 to 1.83 for biochanin A, 0.58 to 1.16 for 2’-hydroxyformononetin, 0.45 to 1.66 for 2’-methoxyformononetin, 0.25 to 2.0 for salicylic acid, 1.08 to 2.83 for jasmonic acid, −0.08 to 1.66 for medicarpin and −2.45 to 0.66 for genistein.

Inter-assay % RE was in the range of −0.33 to 0.50 for daidzein, 0.25 to 1.83 for liquiritigenin, −0.41 to 0.50 for formononetin, −1.16 to1.33 for naringenin, −0.04 to1.50 for biochanin A, 0.25 to 1.33 for 2’-hydroxyformononetin, 0.20 to 1.66 for 2’-methoxyformononetin, −0.41 to 1.00 for salicylic acid, 0.41 to1.00 for jasmonic acid, 0.66 to 0.91 for medicarpin and 0.45 to 2.33 for genistein.

Intra-assay precision (% RSD) was in the range of 0.11 to 0.61 for daidzein, 0.36 to 0.69 for liquiritigenin, 0.46 to 2.06 for formononetin, 0.61 to 1.03 for naringenin, 0.11 to 0.61 for biochanin A, 0.40 to 0.61 for 2’-hydroxyformononetin, 0.31 to 1.01 for 2’-methoxyformononetin, 0.23 to 1.94 for salicylic acid, 0.46 to 1.21 for jasmonic acid, not more than 0.23 for medicarpin and 1.52 to 2.17 for genistein.

The inter-assay % RSD was in the range of 0.10 to 2.06 for daidzein, 0.44 to 0.50 for liquiritigenin, −1.44 to 2.14 for formononetin, 0.41 to 1.26 for naringenin, 0.41 to 0.80 for biochanin A, 0.26 to 0.61 for 2’-hydroxyformononetin, 0.23 to 1.52 for 2’-methoxyformononetin, 0.15 to 1.21 for salicylic acid, 0.35 to 0.93 for jasmonic acid, 0.40 to 2.30 for medicarpin and 0.23 to 1.46 for genistein **(Table 4)**.

**Table 4:** Intraday & Interday Precision (RSD %) and Accuracy (RE %)

| Analyte | Actual Concentration (ng/ml) | Detected concentration (Mean ± SD, n=6) | | RSD (%) | | RE (%) | |
| --- | --- | --- | --- | --- | --- | --- | --- |
|  |  | Intraday | Interday | Intraday | Interday | Intraday | Interday |
| Daidzein | 40.0 | 40.73 ± 0.24 | 39.86 ± 0.82 | 0.61 | 2.06 | 1.83 | -0.33 |
| | 80.0 | 80.20 ± 0.43 | 79.86 ± 0.24 | 0.53 | 0.31 | 0.25 | -0.16 |
| | 160.0 | 160.53 ± 0.18 | 160.8 ± 0.16 | 0.11 | 0.10 | 0.33 | 0.50 |
| Liquiritigenin | 40.0 | 40.8 ± 0.28 | 40.73 ± 0.18 | 0.69 | 0.46 | 2.00 | 1.83 |
| | 80.0 | 81.0 ± 0.56 | 80.93 ± 0.41 | 0.69 | 0.50 | 1.25 | 1.16 |
| | 160.0 | 160.2 ± 0.58 | 160.4 ± 0.71 | 0.36 | 0.44 | 0.12 | 0.25 |
| Formononetin | 40.0 | 39.13 ± 0.18 | 40.20 ± 0.86 | 0.48 | 2.14 | -2.16 | 0.50 |
| | 80.0 | 79.33 ± 1.63 | 80.33 ± 1.15 | 2.06 | -1.44 | -0.83 | 0.41 |
| | 160.0 | 159.46 ± 0.73 | 159.33 ± 0.09 | 0.46 | 0.05 | -0.33 | -0.41 |
| Naringenin | 40.0 | 40.53 ± 0.24 | 39.53 ± 0.49 | 0.61 | 1.26 | 1.33 | -1.16 |
| | 80.0 | 80.93 ± 0.83 | 81.06 ± 0.33 | 1.03 | 0.41 | 1.16 | 1.33 |
| | 160.0 | 161.73 ± 1.55 | 161.06 ± 1.38 | 0.95 | 0.86 | 1.08 | 0.66 |
| Biochanin A | 40.0 | 40.73 ± 0.24 | 40.60 ± 0.32 | 0.61 | 0.80 | 1.83 | 1.50 |
| | 80.0 | 80.46 ± 0.49 | 80.2 ± 0.57 | 0.61 | 0.71 | 0.58 | 0.33 |
| | 160.0 | 160.53 ± 0.18 | 159.93 ± 0.65 | 0.11 | 0.41 | 0.33 | -0.04 |
| 2'-hydroxy formononetin | 40.0 | 40.46 ± 0.18 | 40.53 ± 0.24 | 0.46 | 0.61 | 1.16 | 1.33 |
| | 80.0 | 80.46 ± 0.49 | 80.53 ± 0.24 | 0.61 | 0.30 | 0.58 | 0.66 |
| | 160.0 | 161.46 ± 0.65 | 160.40 ± 0.43 | 0.40 | 0.26 | 0.91 | 0.25 |
| 2'-methoxy formononetin | 40.0 | 40.66 ± 0.41 | 40.66 ± 0.61 | 1.01 | 1.52 | 1.66 | 1.66 |
| | 80.0 | 80.60 ± 0.74 | 80.80 ± 0.43 | 0.92 | 0.53 | 0.75 | 1.00 |
| | 160.0 | 160.73 ± 0.49 | 160.33 ± 0.37 | 0.31 | 0.23 | 0.45 | 0.20 |
| Salicylic acid | 40.0 | 40.80 ± 0.80 | 40.40 ± 0.48 | 1.97 | 1.21 | 2.00 | 1.00 |
| | 80.0 | 80.20 ± 0.28 | 79.66 ± 0.18 | 0.35 | 0.23 | 0.25 | -0.41 |
| | 160.0 | 160.46 ± 0.37 | 160.26 ± 0.24 | 0.23 | 0.15 | 0.29 | 0.16 |
| Jasmonic acid | 40.0 | 41.13 ± 0.49 | 40.33 ± 0.37 | 1.21 | 0.93 | 2.83 | 0.83 |
| | 80.0 | 80.86 ± 0.37 | 80.80 ± 0.28 | 0.46 | 0.35 | 1.08 | 1.00 |
| | 160.0 | 161.73 ± 1.73 | 160.66 ± 0.61 | 1.07 | 0.38 | 1.08 | 0.41 |
| Medicarpin | 40.0 | 40.66 ± 0.09 | 40.26 ± 0.92 | 0.23 | 2.30 | 1.66 | 0.66 |
| | 80.0 | 79.93 ± 0.18 | 80.73 ± 0.67 | 0.23 | 0.84 | -0.08 | 0.91 |
| | 160.0 | 160.46 ± 0.37 | 161.13 ± 0.65 | 0.23 | 0.40 | 0.29 | 0.70 |
| Genistein | 40.0 | 39.86 ± 1.04 | 40.93 ± 0.09 | 2.63 | 0.23 | -0.33 | 2.33 |
|  | 80.0 | 80.53 ± 1.22 | 81.06 ± 0.65 | 1.52 | 0.81 | 0.66 | 1.33 |
|  | 160.0 | 156.06 ± 3.39 | 160.73 ± 2.36 | 2.17 | 1.46 | -2.45 | 0.45 |

### 4.4 Quantification of the flavonoids and phytohormones in *Medicago truncatula*

In this study, a highly sensitive and robust UHPLC-QTRAP-MS/MS method with high reproducibility, accuracy and a short separation time (12 min.) was developed. The method can be used for the quantification of all the metabolites as well as individual molecules for routine analysis. This method, for the simultaneous determination of daidzein, liquiritigenin, formononetin, naringenin, biochanin A, 2’-hydroxy formononetin, 2’-methoxy formononetin, salicylic acid, jasmonic acid, medicarpin and genistein by UHPLC-QTRAP-MS, has not been reported prior to this investigation. Multiple Reaction Monitoring (MRM) was used to quantify these compounds which was based on measuring the parent ion/product ion transitions. The major ion pairs observed in the positive and negative modes were 257/137 for liquiritigenin, 273/153 for naringenin, 269/133 for genistein, 255/199 for daidzein, 269/197 for formononetin, 285/213 for biochanin A, 285/225 for 2’-hydroxy formononetin, 299/148 for 2’-methoxyformononetin, 269/254 for medicarpin, 136.9/65 for salicylic acid and 208.9/59 for jasmonic acid **(Table 1)**. Eleven molecules used as reference standards in the present study showed chromatographic peaks in the total ion chromatogram (TIC) **(Fig. 2A & 2B)** at 2.9, 3.1, 4.7, 3.9, 5.9, 4.1, 4.5, 3.0, 4.1, 4.1 and 3.9 minute for daidzein, liquiritigenin, formononetin, naringenin, biochanin A, 2’-hydroxy formononetin, 2’-methoxyformononetin, salicylic acid, jasmonic acid, medicarpin and genistein respectively.

**Fig. 2B:**
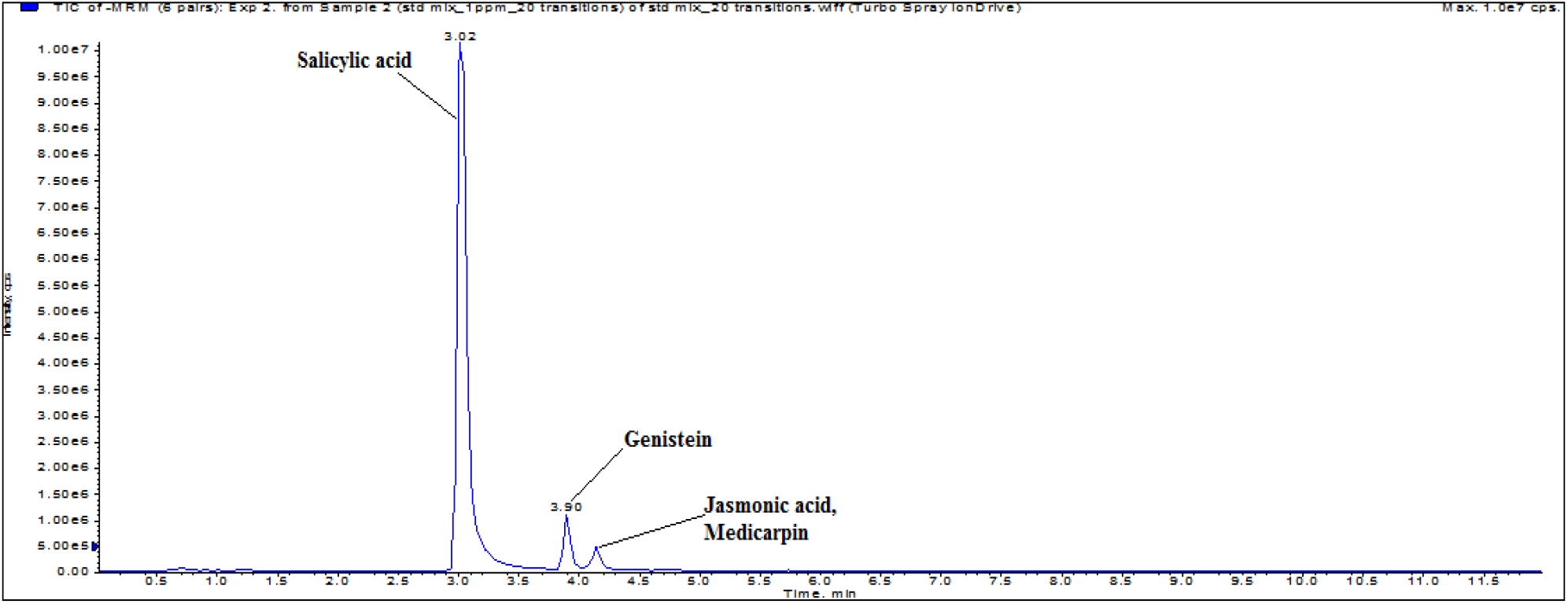
TIC of MRM scan in the negative mode

The current method was successfully employed for the simultaneous identification and quantification of daidzein, liquiritigenin, formononetin, naringenin, biochanin A, 2’-hydroxy formononetin, 2’-methoxy formononetin, salicylic acid, jasmonic acid, medicarpin and genistein in leaf extracts of *M. truncatula.* All eleven metabolites were quantified in leaf extract of *M. truncatula* in triplicates (technical replicates) and values are presented in **Table 5**. We found that formononetin is the most abundant constituent in leaf extract with an average concentration of 5343 ng/ml while biochanin A is the minor constituent with the least average concentration of 30 ng/ml.

**Table 5:** Quantification of metabolites in the leaf extract of *M.truncatula*.

| Analyte | Leaf extract -1_1<br>(ng/ml) | Leaf extract -1_2<br>(ng/ml) | Leaf extract -1_3<br>(ng/ml) | Average<br>concentration<br>(ng/ml) |
| --- | --- | --- | --- | --- |
| Daidzein | 48.0 | 51.0 | 55.5 | 52 |
| Liquiritigenin | 500 | 482 | 480 | 487 |
| Formononetin | 5487 | 5162 | 5381 | 5343 |
| Naringenin | 175 | 177 | 178 | 177 |
| Biochanin A | 37.1 | 33.7 | 18.0 | 30 |
| 2'-hydroxy<br>formononetin | 38.2 | 33.0 | 35.6 | 36 |
| 2'-methoxy<br>formononetin | 53.1 | 54.6 | 54.0 | 54 |
| Salicylic acid | 100 | 113 | 116 | 110 |
| Jasmonic acid | 146 | 173 | 149 | 156 |
| Medicarpin | 68.4 | 68.7 | 87.0 | 75 |
| Genistein | 118 | 149 | 131 | 133 |

#### Concluding remarks

Using UHPLC-QTRAP-MS/MS, a method was developed for the simultaneous detection and quantification of nine flavonoids and two phytohormones and has been successfully applied for the quantitation of these metabolites in the leaf extract of *Medicago Truncatula*. Satisfactory validation parameters were obtained including linearity, specificity, accuracy, precision and stability. The same method can be applied for the quantitation of these metabolites in the other plant products and assays.

## Acknowledgements

This research work was carried out at the Mass spectrometry facility of the Advanced Technology Platform Centre (ATPC), managed by the Regional Centre for Biotechnology (RCB), Faridabad (HRY)-India. The authors NS and NSh are highly thankful to the **Department of Biotechnology (**Grant No. BT.MED-II/ ATPC/BSC/01/2010) for their financial support to the facility. PA acknowledges fellowship grants from the Council of Scientific and Industrial Research.

