## Supplementary data for "Development of a rapid LC-MS/MS method for the simultaneous quantification of various flavonoids and phytohormones extracted from *Medicago Truncatula* leaves"

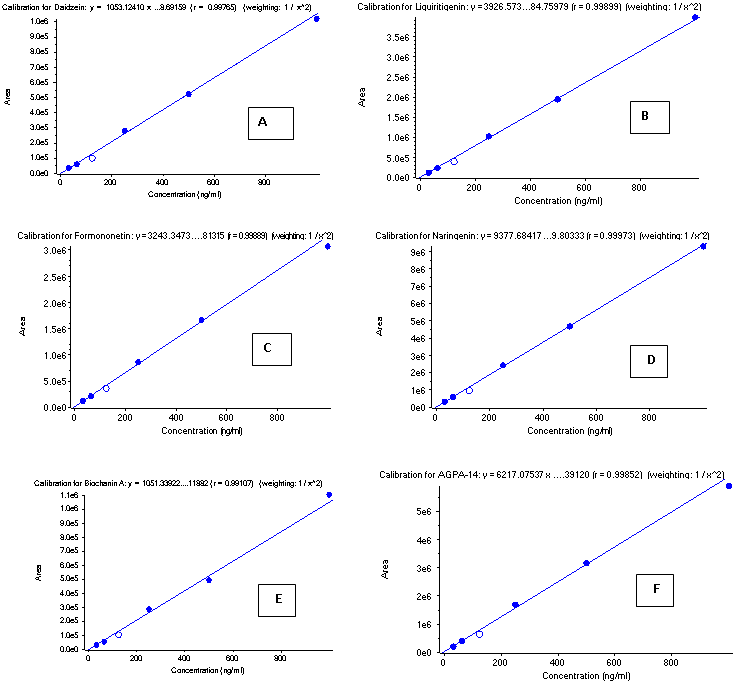

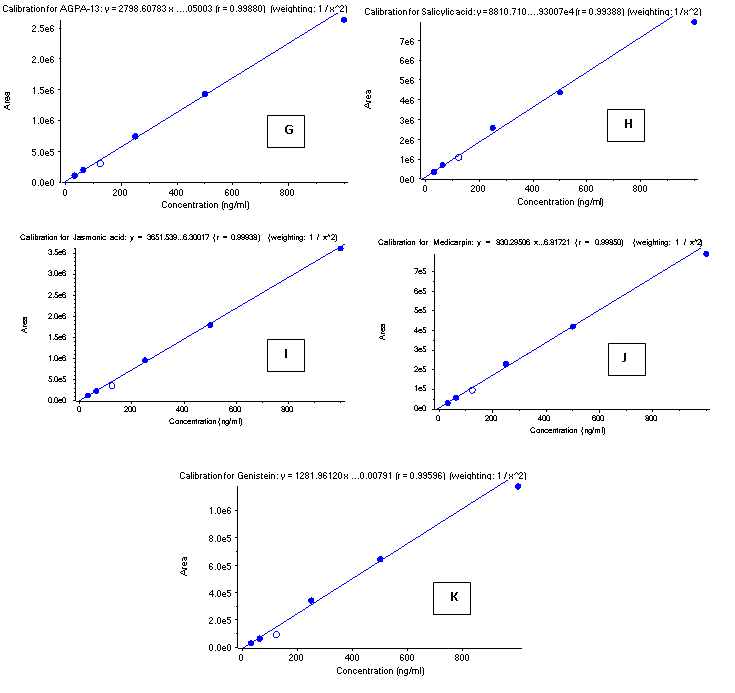

**Fig. S1:** Six-point calibration curves of (A) Daidzein (B) Liquiritigenin (C) Formononetin (D)Naringenin (E) Biochanin A (F) 2'-hydroxy formononetin (G) 2'-methoxy formononetin (H) Salicylic acid (I) Jasmonic acid (J) Medicarpin (K) Genistein

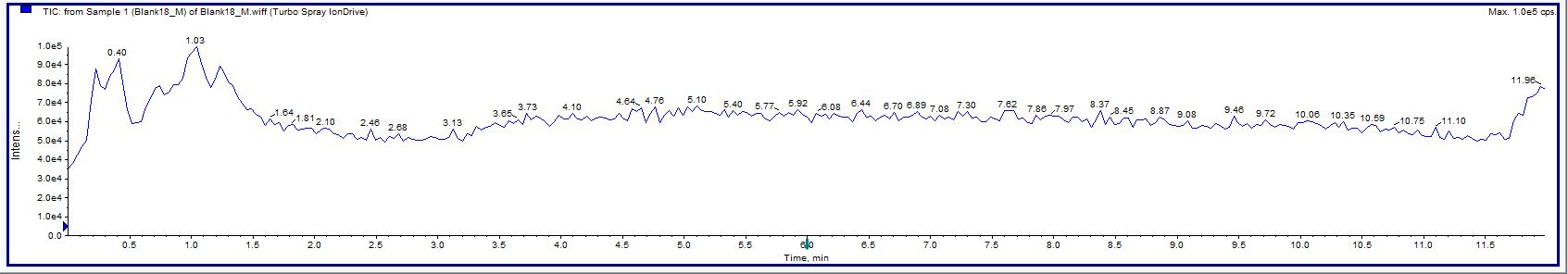

**Fig. S2:** Total ion chromatogram of Blank

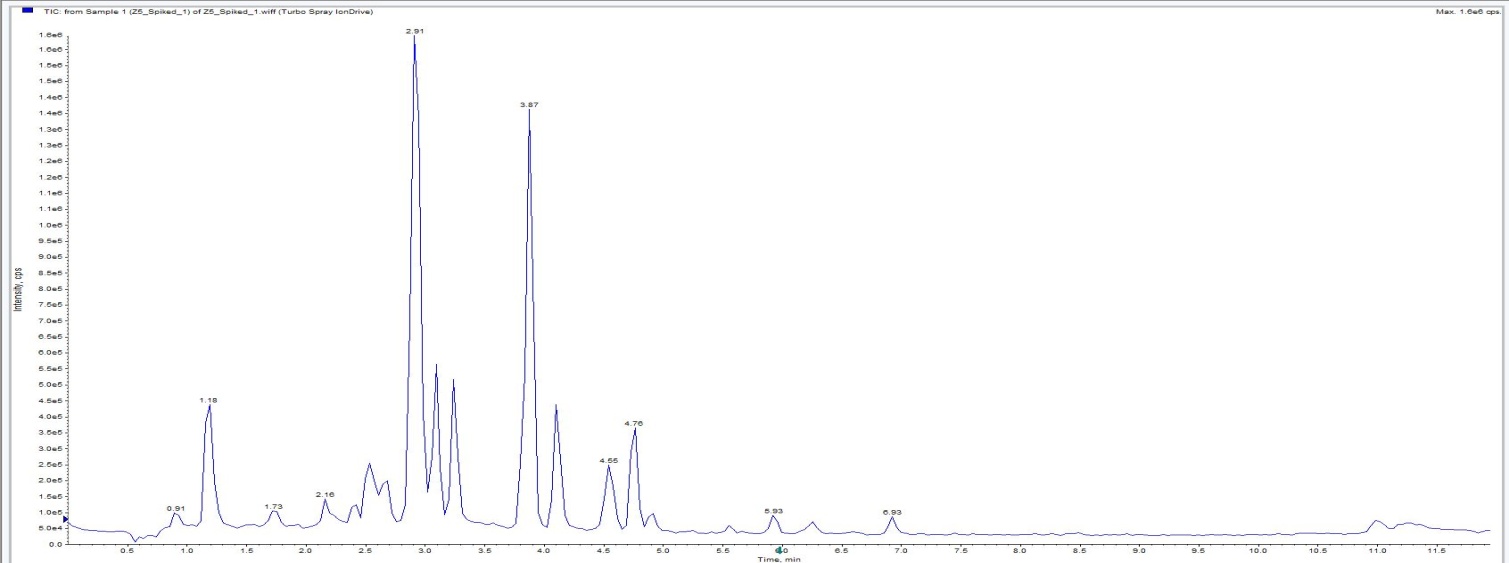

**Fig. S3:** Total ion Chromatogram of extract spiked with standards (low level spiking at 10ng/ml)
